## Supplemental Materials for "Modeling functional cell types in spike train data"

### Supplement

February 27, 2023

#### S1 Extended EM Methods

Here we provide a more detailed account of how our adapted EM algorithm fits our hierarchical generative model (11) to data.

##### S1.1 E-Step

Our expectation (E) step consists of finding the $Z_{i,k}, \boldsymbol{m}_{i,k}, c_{i,k}$ that best approximate the posterior distribution over the latent variables, $Q_i(k, \boldsymbol{\beta})$ (17).

For fixed $i$ and $k$, we first set $\boldsymbol{m}_{i,k}$ so that the modes of the left and right hand sides of (14) are equal. To do this, we apply the trust-region Newton-conjugate-gradient algorithm to solve the convex optimization problem

$$\boldsymbol{m}_{i,k} = \arg\max_{\boldsymbol{\beta}} \log P_{\text{joint}}(k, \boldsymbol{\beta}, \mathbf{y}_i | \mathbf{x}_i; \hat{\Omega}_K). \tag{S1}$$

Next, so that the Hessians of the logs of the left and right hand sides of (14) are equal, we compute

$$\tilde{c}_{i,k}^{-1} = -\nabla^2_{\boldsymbol{\beta}} \log P_{\text{joint}}(k, \boldsymbol{\beta}, \mathbf{y}_i | \mathbf{x}_i; \hat{\Omega}_K)\bigg|_{\boldsymbol{\beta}=\boldsymbol{m}_{i,k}}. \tag{S2}$$

The Hessian of a GLM log-likelihood function is often used during optimization (see [1] 9.3.2 for details), and we make use of the `statsmodels` package to efficiently compute it. We further approximate $\tilde{c}_{i,k}^{-1}$ by retaining only its diagonal elements, i.e. $c_{i,k}^{-1} = \text{diag}(\tilde{c}_{i,k})^{-1}$.

Finally, noting that $f(\boldsymbol{\beta}; \boldsymbol{m}_{i,k}, c_{i,k})\Big|_{\boldsymbol{\beta}=\boldsymbol{m}_{i,k}} = \frac{1}{\sqrt{(2\pi)^{T^{\text{stim}}+T^{\text{self}}+1}|c_{i,k}|}}$, we take

$$Z_{i,k} = P_{\text{joint}}(k, \boldsymbol{\beta}, \mathbf{y}_i | \mathbf{x}_i; \hat{\Omega}_K)\bigg|_{\boldsymbol{\beta}=\boldsymbol{m}_{i,k}} \sqrt{(2\pi)^{T^{\text{stim}}+T^{\text{self}}+1}|c_{i,k}|}, \tag{S3}$$

where $|c_{i,k}|$ denotes the determinant, so that the left and right hand sides of (14) are equal when $\boldsymbol{\beta} = \boldsymbol{m}_{i,k}$. Collectively, (S1), (S2), and (S3) are known as the Laplace approximation (see [1] 8.4.1).

This concludes the set of operations needed to perform the E-step.

##### S1.2 M-Step

The maximization (M) step consists of finding $\Omega_K$ that maximizes the lower bound on the log-likelihood, (18). In light of (8), the bound being maximized in the M-step factors as follows:

$$\sum_{i=1}^{N}\sum_{k=1}^{K}\int \tilde{Z}_{i,k} f(\boldsymbol{\beta};\boldsymbol{m}_{i,k},c_{i,k}) \log P_{\text{joint}}(k,\boldsymbol{\beta},\mathbf{y}_i|\mathbf{x}_i;\Omega_K) d\boldsymbol{\beta}$$

$$= \sum_{i=1}^{N}\sum_{k=1}^{K}\int \tilde{Z}_{i,k} f(\boldsymbol{\beta};\boldsymbol{m}_{i,k},c_{i,k}) [\log P_{\text{SC}}(\mathbf{y}_i|\mathbf{x}_i,\boldsymbol{\beta}) + \log \pi_k + \log f(\boldsymbol{\beta};\boldsymbol{\mu}_k,\Sigma_k)] d\boldsymbol{\beta}.$$

Absorbing into $C$ terms that do not depend on the optimization variables $\Omega_K$, this equa[illegible]

$$\sum_{i=1}^{N}\sum_{k=1}^{K}\int \tilde{Z}_{i,k} f(\boldsymbol{\beta};\boldsymbol{m}_{i,k},c_{i,k}) [\log \pi_k + \log f(\boldsymbol{\beta};\boldsymbol{\mu}_k,\Sigma_k)] d\boldsymbol{\beta} + C.$$

Rearranging the sums and integrals, this equals

$$\sum_{k=1}^{K}\int [\sum_{i=1}^{N} \tilde{Z}_{i,k} f(\boldsymbol{\beta};\boldsymbol{m}_{i,k},c_{i,k})] [\log \pi_k + \log f(\boldsymbol{\beta};\boldsymbol{\mu}_k,\Sigma_k)] d\boldsymbol{\beta} + C.$$

Distributing the product and using $\int f(\boldsymbol{\beta};\boldsymbol{m}_{i,k},c_{i,k}) d\boldsymbol{\beta} = 1$ to simplify the first term, th[illegible]

$$\sum_{k=1}^{K}[\sum_{i=1}^{N} \tilde{Z}_{i,k}] \log \pi_k + \sum_{k=1}^{K}\int [\sum_{i=1}^{N} \tilde{Z}_{i,k} f(\boldsymbol{\beta};\boldsymbol{m}_{i,k},c_{i,k})] \log f(\boldsymbol{\beta};\boldsymbol{\mu}_k,\Sigma_k) d\boldsymbol{\beta} + C.$$

We maximize the first term in (S4), $\sum_{k=1}^{K}[\sum_{i=1}^{N} \tilde{Z}_{i,k}] \log \pi_k$, with respect to $\pi_1,...,\pi_K$, subject to $\pi_k \geq 0, k = 1,...,K$ and $\sum_{k=1}^{K} \pi_k = 1$. Algebraic manipulation reveals that

$$\hat{\pi}_k = \frac{1}{N}\sum_{i=1}^{N} \tilde{Z}_{i,k}. \tag{S5}$$

Each summand in the second term in (S4), $\int [\sum_{i=1}^{N} \tilde{Z}_{i,k} f(\boldsymbol{\beta};\boldsymbol{m}_{i,k},c_{i,k})] \log f(\boldsymbol{\beta};\boldsymbol{\mu}_k,\Sigma_k) d\boldsymbol{\beta}$, is maximized independently with respect to $\boldsymbol{\mu}_k$ and $\Sigma_k$. We use the probability density function for a multivariate Gaussian to rewrite this term as

$$\int \left[\sum_{i=1}^{N} \tilde{Z}_{i,k} f(\boldsymbol{\beta};\boldsymbol{m}_{i,k},c_{i,k})\right] \left[-\frac{(\boldsymbol{\beta}-\boldsymbol{\mu}_k)^\top \Sigma_k^{-1}(\boldsymbol{\beta}-\boldsymbol{\mu}_k) - (T^{\text{stim}} + T^{\text{self}} + 1)\log(2\pi) + \log(|\Sigma_k^{-1}\text{[illegible]}}{2}\right. \tag{S6}$$

Differentiating (S6) with respect to $\boldsymbol{\mu}_k$ reveals that

$$\begin{aligned} 0 &= \int \left[\sum_{i=1}^{N} \tilde{Z}_{i,k} f(\boldsymbol{\beta};\boldsymbol{m}_{i,k},c_{i,k})\right] \Sigma_k^{-1}(\boldsymbol{\beta}-\boldsymbol{\mu}_k) d\boldsymbol{\beta} \\ &= \sum_{i=1}^{N} \tilde{Z}_{i,k}\Sigma_k^{-1}\left[\int f(\boldsymbol{\beta};\boldsymbol{m}_{i,k},c_{i,k})\boldsymbol{\beta} d\boldsymbol{\beta} - \int f(\boldsymbol{\beta};\boldsymbol{m}_{i,k},c_{i,k})\boldsymbol{\mu}_k d\boldsymbol{\beta}\right] \\ &= \sum_{i=1}^{N} \tilde{Z}_{i,k}\Sigma_k^{-1}\left[\boldsymbol{m}_{i,k} - \boldsymbol{\mu}_k\right]. \end{aligned}$$

This gives us

$$\hat{\boldsymbol{\mu}}_k = \frac{\sum_{i=1}^{N} \tilde{Z}_{i,k}\boldsymbol{m}_{i,k}}{\sum_{i=1}^{N} \tilde{Z}_{i,k}}. \tag{S7}$$

To maximize (S6) with respect to $\Sigma_k$, we note that it is concave in $\Sigma_k^{-1}$. We can then solve for $\Sigma_k$ by setting the derivative with respect to $\Sigma_k^{-1}$ equal to zero:

$$0 = \int \sum_{i=1}^{N} \tilde{Z}_{i,k} f(\boldsymbol{\beta}; \boldsymbol{m}_{i,k}, c_{i,k}) \left[ -\frac{1}{2}(\boldsymbol{\beta} - \boldsymbol{\mu}_k)(\boldsymbol{\beta} - \boldsymbol{\mu}_k)^\top + \frac{1}{2}\Sigma_k \right] d\boldsymbol{\beta},$$

making use of the fact that $\frac{d}{dX}\log(|X|) = (X^{-1})^\top$, and that $\Sigma_k = \Sigma_k^\top$. That is,

$$\Sigma_k = \frac{\sum_{i=1}^{N} \tilde{Z}_{i,k} \mathbb{E}\left[(\boldsymbol{\beta} - \boldsymbol{\mu}_k)(\boldsymbol{\beta} - \boldsymbol{\mu}_k)^\top\right]}{\sum_{i=1}^{N} \tilde{Z}_{i,k}},$$

where the expectation is with respect to $\boldsymbol{\beta} \sim \mathcal{N}(\boldsymbol{m}_{i,k}, c_{i,k})$. We can decompose this ex[illegible]

$$\begin{aligned}
&\mathbb{E}_{\boldsymbol{\beta}\sim\mathcal{N}(\boldsymbol{m}_{i,k}, c_{i,k})}\left[(\boldsymbol{\beta} - \boldsymbol{\mu}_k)(\boldsymbol{\beta} - \boldsymbol{\mu}_k)^\top\right] \\
= &\mathbb{E}_{\boldsymbol{\beta}\sim\mathcal{N}(\boldsymbol{m}_{i,k}, c_{i,k})}\left[(\boldsymbol{\beta} - \boldsymbol{m}_{i,k} + \boldsymbol{m}_{i,k} - \boldsymbol{\mu}_k)(\boldsymbol{\beta} - \boldsymbol{m}_{i,k} + \boldsymbol{m}_{i,k} - \boldsymbol{\mu}_k)^\top\right] \\
= &\mathbb{E}_{\boldsymbol{\beta}\sim\mathcal{N}(\boldsymbol{m}_{i,k}, c_{i,k})}\left[(\boldsymbol{\beta} - \boldsymbol{m}_{i,k})(\boldsymbol{\beta} - \boldsymbol{m}_{i,k})^\top + (\boldsymbol{m}_{i,k} - \boldsymbol{\mu}_k)(\boldsymbol{m}_{i,k} - \boldsymbol{\mu}_k)^\top \right.\\
&\left. + (\boldsymbol{\beta} - \boldsymbol{m}_{i,k})(\boldsymbol{m}_{i,k} - \boldsymbol{\mu}_k)^\top + (\boldsymbol{m}_{i,k} - \boldsymbol{\mu}_k)(\boldsymbol{\beta} - \boldsymbol{m}_{i,k})^\top\right] \\
= &c_{i,k} + (\boldsymbol{m}_{i,k} - \boldsymbol{\mu}_k)(\boldsymbol{m}_{i,k} - \boldsymbol{\mu}_k)^\top.
\end{aligned}$$

Plugging this expression back in for the expectation, we have

$$\begin{aligned}
\Sigma_k &= \frac{\sum_{i=1}^{N} \tilde{Z}_{i,k}(c_{i,k} + \boldsymbol{m}_{i,k}\boldsymbol{m}_{i,k}^\top + \boldsymbol{\mu}_k\boldsymbol{\mu}_k^\top - \boldsymbol{m}_{i,k}\boldsymbol{\mu}_k^\top - \boldsymbol{\mu}_k\boldsymbol{m}_{i,k}^\top)}{\sum_{i=1}^{N} \tilde{Z}_{i,k}} \\
&= \frac{\sum_{i=1}^{N} \tilde{Z}_{i,k}(c_{i,k} + \boldsymbol{m}_{i,k}\boldsymbol{m}_{i,k}^\top - \boldsymbol{\mu}_k\boldsymbol{\mu}_k^\top)}{\sum_{i=1}^{N} \tilde{Z}_{i,k}},
\end{aligned}$$

where in the last step we make use of (S7) to simplify the last two terms in the numerator. Finally, subjecting this expression to the constraint that $\Sigma_k$ must be diagonal, we have:

$$\hat{\Sigma}_k = \frac{\sum_{i=1}^{N} \tilde{Z}_{i,k} \mathrm{diag}(c_{i,k} + \boldsymbol{m}_{i,k}\boldsymbol{m}_{i,k}^\top - \hat{\boldsymbol{\mu}}_k\hat{\boldsymbol{\mu}}_k^\top)}{\sum_{i=1}^{N} \tilde{Z}_{i,k}}. \tag{S8}$$

In the main manuscript, the solutions (S7) and (S8) are applied only to $\boldsymbol{\mu}^{\mathrm{self}}, \Sigma^{\mathrm{self}}$, as the other components are held fixed during EM.

#### S2 Alternative Model: all parameters depend on cell-type

Here, we consider an alternate formulation of both sequential and simultaneous methods, where instead of just $\boldsymbol{\beta}_i^{\mathrm{self}}$ being related to cell-type, as considered in the main manuscript, we consider the case where all of $\boldsymbol{\beta}_i$ is related to cell-type. We call the former, in main, case A, and the latter, detailed here, case B.

##### S2.1 Alternative Methods

Below we detail how the sequential and simultaneous methods change for case B. For both, the main difference is that instead of fitting a cluster model with means $\boldsymbol{\mu}_k^{\mathrm{self}}$ and covariance matrices $\Sigma_k^{\mathrm{self}}$ that describes the distribution of $\boldsymbol{\beta}_i^{\mathrm{self}}$, they now fit one with means $\boldsymbol{\mu}_k$ and covariance matrices $\Sigma_k$ that describes the distribution of $\boldsymbol{\beta}_i$. For convenience, we will refer to the appropriate components of $\boldsymbol{\mu}_k$ and $\Sigma_k$ using superscripts, e.g. $\boldsymbol{\mu}_k^{\mathrm{stim}}$, just as we do with $\boldsymbol{\beta}_i$.

###### S2.1.1 Sequential Method

Because, in the sequential method, the fitting of single cell models does not depend on cell-types, that first stage is identical between cases A and B. For the second stage, clustering, the only difference is that GMM clustering is performed on all of $\hat{\boldsymbol{\beta}}_i$, instead of just $\hat{\boldsymbol{\beta}}_i^{\text{self}}$:

$$\hat{\Omega}_K \leftarrow \arg\max_{\Omega_K} \sum_{i=1}^{N} \log \sum_{k=1}^{K} \pi_k f(\hat{\boldsymbol{\beta}}_i; \boldsymbol{\mu}_k, \Sigma_k) \tag{S9}$$

$$\hat{k}_i \leftarrow \arg\max_k \hat{\pi}_k f(\hat{\boldsymbol{\beta}}_i; \hat{\boldsymbol{\mu}}_k, \hat{\Sigma}_k),\ i = 1, ..., N \tag{S10}$$

Algorithm 1 is thus unchanged except for changing the last two lines accordingly. The computation of the BIC also changes accordingly:

$$BIC \equiv \sum_{i=1}^{N} \log \left[ \sum_{k=1}^{K} \hat{\pi}_k f(\hat{\boldsymbol{\beta}}_i; \hat{\boldsymbol{\mu}}_k, \hat{\Sigma}_k) \right] - \frac{K - 1 + 2K\text{dim}(\boldsymbol{\beta})}{2} \log(N) \tag{S11}$$

###### S2.1.2 Simultaneous Method

For the simultaneous method, case B is much simpler, as all parameters of the single cell models use the learned cluster structure as priors, so there is no longer a need for $\ell_2$ regularization.

The joint probability of the data and parameters simplifies instead to:

$$P_{\text{joint}}(k, \boldsymbol{\beta}, \mathbf{y}_i | \mathbf{x}_i; \Omega_K) \propto P_{SC}(\mathbf{y}_i | \mathbf{x}_i; \boldsymbol{\beta}) f(\boldsymbol{\beta}_i^{\text{stim}}; \boldsymbol{\mu}_k^{\text{stim}}, \Sigma_k^{\text{stim}}) f(\boldsymbol{\beta}_i^{\text{self}}; \boldsymbol{\mu}_k^{\text{self}}, \Sigma_k^{\text{self}}) f(\beta_i^0; \mu_k^0, \Sigma_k^0) \pi_k. \tag{S12}$$

With this adapted $P_{\text{joint}}$ the results derived in the main manuscript, i.e. (12)-(21), all apply to case B, without making the exceptions for constrained cluster means and covariances, as was done for S7,S8. Algorithm 2 is thus unchanged except that all of $\boldsymbol{\mu}_k$ and $\Sigma_k$ are updated in the M-step instead of just $\boldsymbol{\mu}_k^{\text{self}}$ and $\Sigma_k^{\text{self}}$. For (21), in case B we have $\text{dof}(\hat{\Omega}_K) = K * (2 * (T^{\text{self}} + T^{\text{stim}} + 1) + 1) - 1$, reflecting the change in number of free parameters.

##### S2.2 Results on Simulated Data

Here we show the equivalent results to those presented in Figures 2 and 3, but for the alternative model where all parameters depend on cell-type. For these results, the simulated dataset was sampled from GLMs whose true $\boldsymbol{\mu}^{\text{stim}}$ and $\mu^0$ were clustered by cell-type, in addition to $\boldsymbol{\mu}^{\text{self}}$ (Figure S1 A,B).

Figure S1C-F show very similar results for parameter recovery to those in Figure 2, the only salient difference being that the recovery of $\boldsymbol{\beta}_i^{\text{stim}}$ is also much better in the simultaneous method, and for lower $\sigma$. This is to be expected now that the true values of these parameters depend on sigma, and their estimates in the simultaneous method make use of cell-type specific priors.

Model selection, however, has markedly worse performance with the all-parameters-shared model, and we can no longer say that the simultaneous method is better (Figure S2). Since the $\boldsymbol{\mu}^{\text{self}}$ are the same in all cases we consider, we can interpret this result by saying that for our choice of $\Omega_K$, the additional differences between clusters in $\boldsymbol{\mu}^{\text{stim}}$ and $\mu^0$ do not make it easier enough to separate the clusters to overcome the increased complexity penalty in BIC with increased $\text{dof}(\hat{\Omega}_K)$.

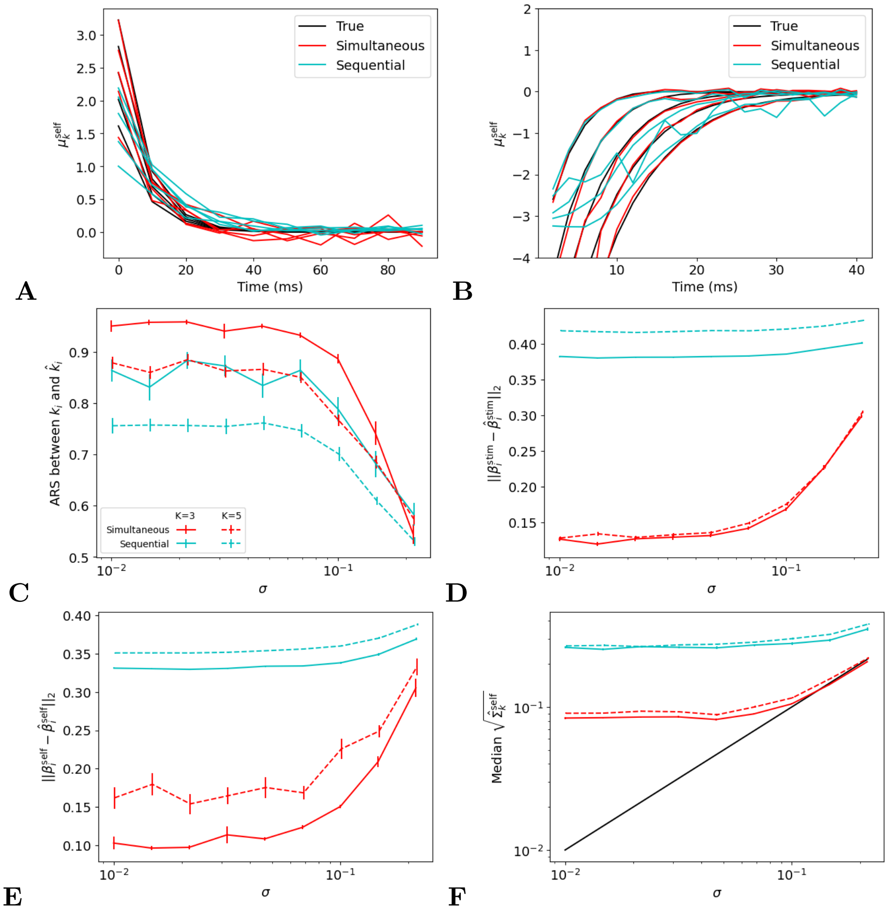


**Fig S1. Performance of simultaneous and sequential methods with all parameters shared on data simulated from a fixed cluster structure.**
A, B: The true cluster means $\boldsymbol{\mu}_k^{\text{stim}}, \boldsymbol{\mu}_k^{\text{self}}$ used to generate simulated datasets and those estimated by the sequential and simultaneous methods, fit with the correct $K = 5$.
C-F: same plots as Figure 2, but with the alternative model. In all metrics and conditions except ARS with $K = 3$ and the two highest values of $\sigma$, the simultaneous method is significantly better.

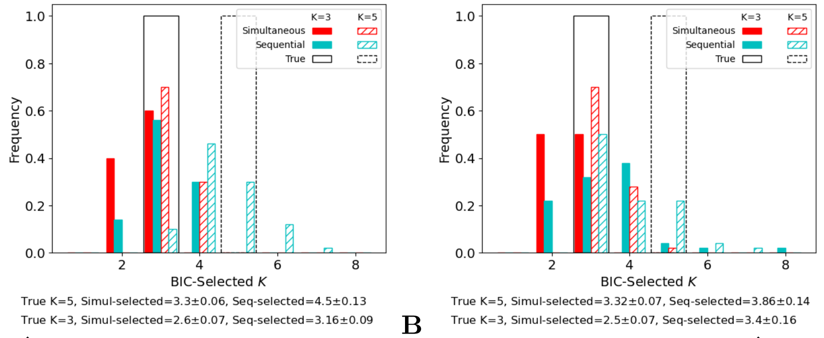


**Fig S2. Model selection of $\hat{K}$ using BIC with the alternative model.** Frequencies of $\hat{K}$ estimated via BIC over 50 simulated datasets with the same $\mu_k$ as in Figure S1, and $\sigma = 10^{-2}$ (A) or $10^{-5/6}$ (B), the maximum value that does not result in degenerate simulations. Black lines indicate true $K$. Summary below plots gives the mean ± SEM of estimated $\hat{K}$ across the 50 datasets for each case and each method.

#### S2.3 Results on IVSCC Data

Here we show results like those shown in Figures 4,5, and 6, but using the alternate model where all parameters are cell-type-dependent (case B).

In Figures S3, S4, and S5, we compare between simultaneous and sequential methods the discovered cluster structure, the generalization performance of fitted models, and the link between cluster labels and metadata, respectively, resulting from fitting to the IVSCC data. The results are comparable to those in the main manuscript.

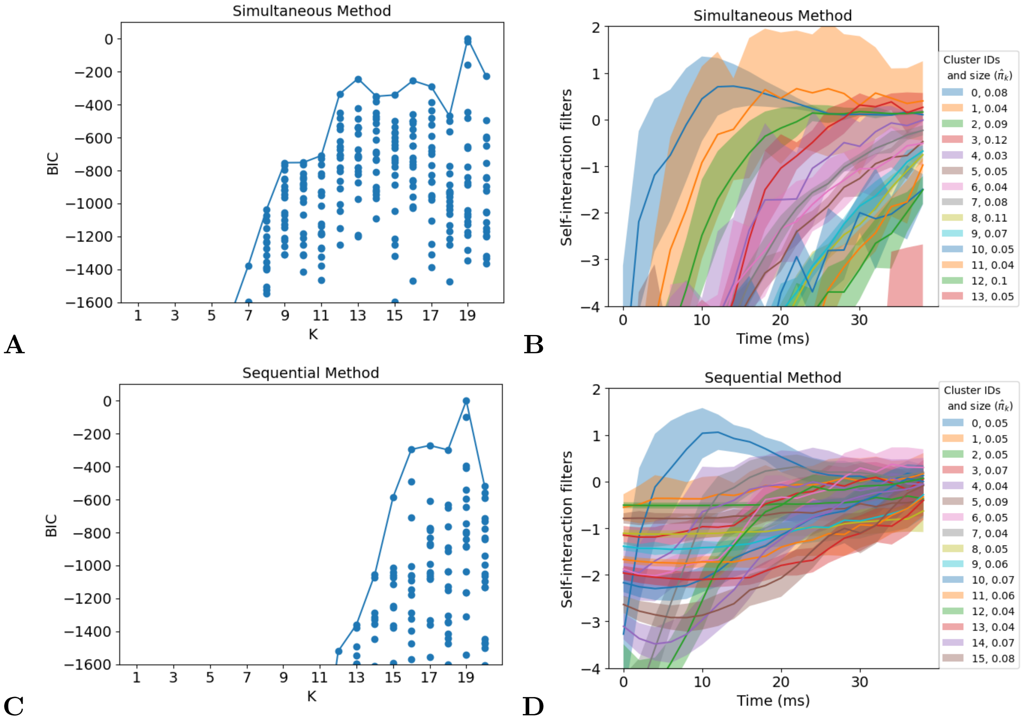


**Fig S3. IVSCC Dataset, Alternative Model.** Same analysis as in Figure 4, but with the alternative model (case B) where all parameters are cell-type dependent. Here, $\hat{K} = 19$ is selected by BIC for both methods.

### S3 Additional details for Figures 5 and S4

The following descriptions apply to both Figures 5 and S4, except for panels B, D, and F, which do not exist in Figure S4, as these analyses have not been performed. The results in Figure 5 are from models where only the self-interaction filters are shared ("Case A" - see Sections 2.2 and 2.3), whereas those in Figure S4 are from models where all parameters are shared ("Case B" - see Sections S2.1.1 and S2.1.2).

To generate panels A, C, and E, we split the neurons into four folds, and applied both simultaneous and sequential approaches and performed model selection on three of the four folds using the Noise 1 stimulus. We then fit GLM models for each neuron in the test fold using the Noise 1 stimulus with the results of training (selected hyperparameters, as well as the fitted hierarchical model for the simultaneous method). Then, we use the Noise 2 stimulus to evaluate those test neurons. We repeat this process for each choice of test fold to evaluate all neurons.

To generate panels B, D, and F:

- We fix all hyperparameters to their values selected when using all presentations of Noise 1 to all neurons.
- We split the neurons into four folds, and applied both simultaneous and sequential

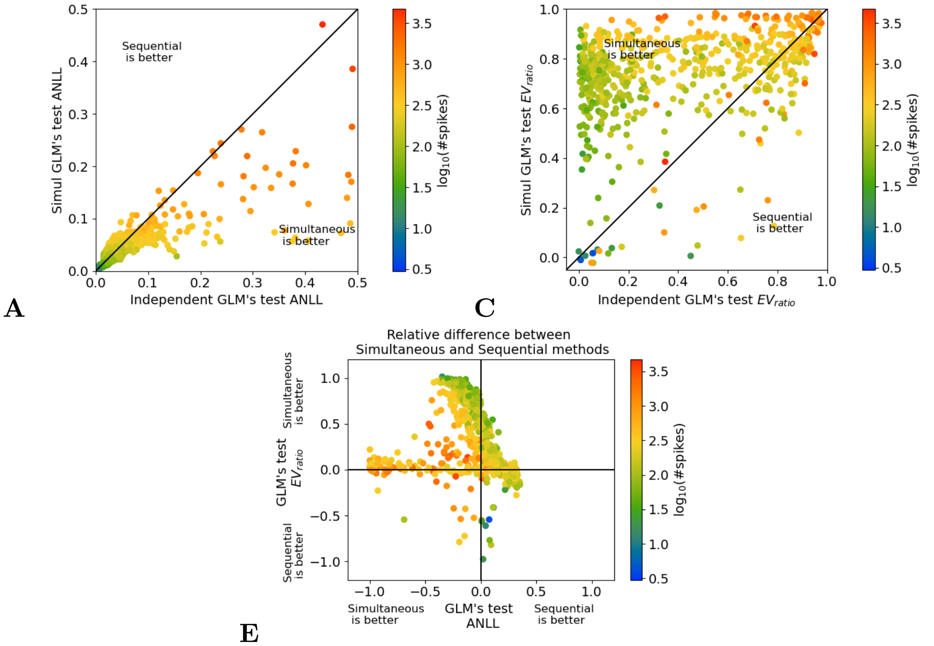


**Fig S4. IVSCC Generalization Performance, Alternative Model.** Same analysis as Figure 5A,C, and E, but with the alternative model (case B) where all parameters are cell-type dependent. Results here are comparable to those in the main manuscript.

approaches to subsets of the folds: either one fold, or two folds, or three folds. These are the columns of panels B, D, and F. We also used only one, two, or three presentations of the Noise 1 stimulus: these are the rows of panels B, D, and F. We then estimate the GLM parameters and cluster assignments for all neurons in the test fold(s), using all presentations of the Noise 1 stimulus with the fixed hyperparameters and the results of training (the fitted hierarchical model for the simultaneous method or the fitted GMM for the sequential method). We then evaluated ANLL and $EV_{ratio}$ on all presentations of Noise 2 to these test neurons. This process is then repeated for each possible division of the folds into training and testing, and ANLL and $EV_{ratio}$ for each neuron are each averaged over all divisions in which that neuron was in the test fold(s).

For instance, to obtain the top left square of panel B, we repeatedly applied both simultaneous and sequential approaches to $1/4$ of the neurons (one fold) using only one presentation of the Noise 1 stimulus. Then, for the remaining $3/4$ of the neurons (three folds), we estimated their GLM parameters using all presentations of Noise 1 (along with the fitted hierarchical model for the simultaneous method), and finally evaluated them using all presentations of the Noise 2 stimulus.

- In each cell of panels B and D, we define a *sample* as the relative difference (simultaneous minus sequential, divided by their sum) in ANLL and $EV_{ratio}$, respectively, of a single neuron.
- In each cell of panel F, we define a *sample* as the relative difference (simultaneous minus sequential, divided by their sum) in ARS of cluster assignments between a pair of different divisions of the neurons into training and testing.
- In each cell, the color is the median across samples, and we place an asterisk if the simultaneous method is significantly better (negative relative difference for panel B, positive for D and F), i.e. if a one-sided Wilcoxon signed-rank test of the samples produces an uncorrected p-value less than 0.001.

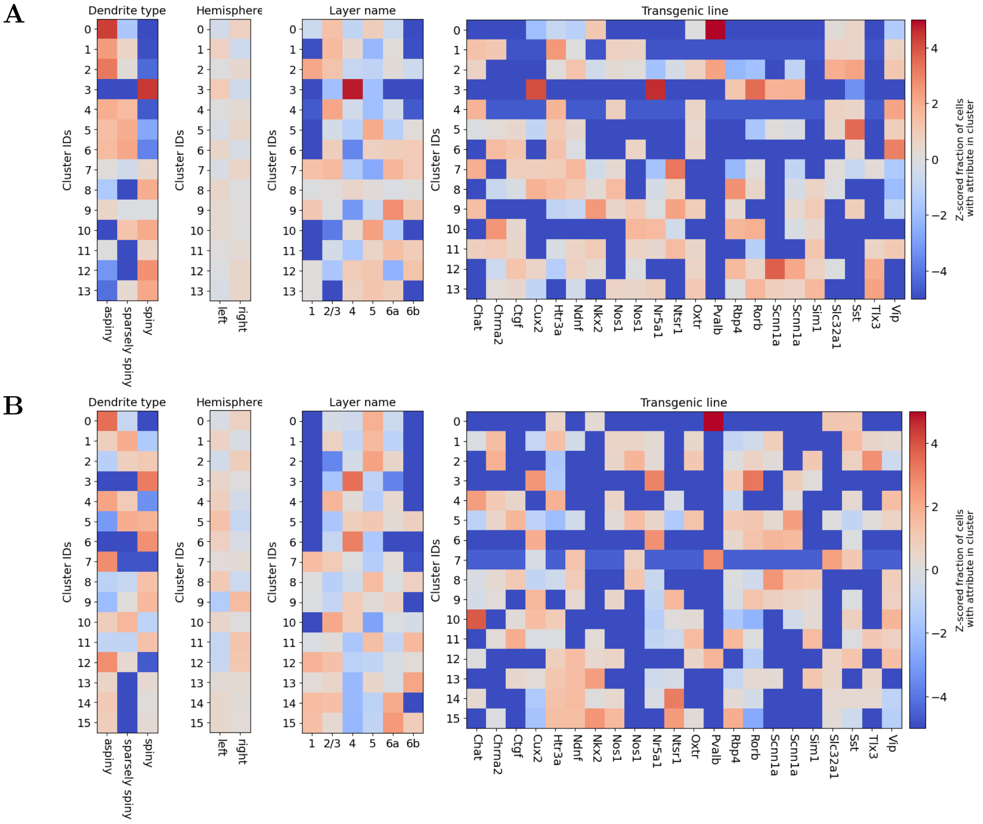


**Fig S5. IVSCC Metadata, alternate model.** Same analysis as Figure 6, but with the alternative model (case B) where all parameters are cell-type dependent. Results here are comparable to those in the main manuscript.

- Between each pair of adjacent cells, except horizontal pairs in panel F, we computed a two-sided Wilcoxon signed-rank test on the paired differences between their samples. We placed a dash if the p-value is less than 0.001.
- The p-values reported in the caption for trends in horizontal and vertical differences, except for the horizontal trend in panel F, are calculated by a one-sided Wilcoxon signed-rank test on the paired differences between samples, aggregated across all six pairs of either horizontally or vertically adjacent cells. P-values greater than 0.1 are reported as "not significant."
- Between each pair of horizontally adjacent cells in panel F, samples cannot be paired, so we used a two-sided, two-sample t-test between the samples from the left and right cells to measure significance, and placed a dash if the p-value is less than 0.001.
- Likewise, the p-value for the horizontal trend in panel F is computed by a one-sided two-sample t-test between the aggregated samples from the six leftmost cells and the six rightmost cells (it was greater than 0.1 and thus labeled "not significant").

#### S4 Parameters used for Simulated Data

This section details our process for simulating datasets from hierarchical models in both cases A and B. These datasets were used to generate Figures 2 and 3 and Figures S1 and S2 respectively.

Instead of truly sampling the cell-types, $k_i$, from $\{\pi_k\}$ we simply assign an equal number (40) of neurons to each class. To set the filters, we use functions that parameterize the stimulus filters as a decaying exponential,

$$g(t; a_{stim}, \tau_{stim}) \equiv a_{stim} e^{-\frac{t}{\tau_{stim}}}, \tag{S13}$$

and the self-interaction filters as the sum of a negative decaying exponential and a Gaussian bump,

$$h(t; t_{ref}, \tau_{ref}, a_{ISI}, \mu_{ISI}, \sigma_{ISI}) \equiv -e^{-\frac{t-t_{ref}}{\tau_{ref}}} + a_{ISI} e^{-\frac{(t-\mu_{ISI})^2}{2*\sigma_{ISI}^2}}. \tag{S14}$$

In both cases A and B, $\boldsymbol{\mu}_k^{\text{self}}$ are determined with $t_{ref}, \tau_{ref}, a_{ISI}, \mu_{ISI}, \sigma_{ISI}$ that are set to uniformly spaced values for each $k$. In case A, $a_{stim}, \tau_{stim}$, and $\beta_k^0$ are each fixed to a single value for all neurons, whereas in case B they are also set to uniformly spaced values for each $\boldsymbol{\mu}_k$. All relevant constants that parameterize this spacing are enumerated in Table 1.

| Evenly spaced params for $\boldsymbol{\mu}_k^{\text{self}}$ in both cases A and B | | | |
|---|---|---|---|
| Parameter | Value for $\boldsymbol{\mu}_1$ | Increment | |
| $t_{ref}$ | 2 | 1.75 | |
| $\tau_{ref}$ | 2 | 0.5 | |
| $a_{ISI}$ | 0.2 | $-0.05$ | |
| $\mu_{ISI}$ | 3 | 2 | |
| $\sigma_{ISI}$ | 3 | 0.25 | |
| Evenly spaced for $\boldsymbol{\mu}_k$ in case B, fixed to one value for all $\boldsymbol{\beta}_i$ in case A | | | |
| Parameter | Value for $\boldsymbol{\mu}_1$ (case B) | Increment (case B) | Value for all $\boldsymbol{\beta}_i$ (case A) |
| $a_{stim}$ | 0.5 | 0.125 | 0.9 |
| $\tau_{stim}$ | 4 | 0 | 4 |
| $\mu_k^0$ | $-4.5$ | 0.25 | -5 |

**Table 1.** Spacing of the parameters used to simulate datasets for cases A and B.

We also used the same stimulus downsampling factor, $d^{\text{stim}} = 5$, in our simulations. In order that its value does not affect the simulation much, we compute $g(t), t \in \{1, .., T^{\text{stim}} * d^{\text{stim}}\}$, and then sum the $\tau$th block of $d^{\text{stim}}$ values, $\sum_{t=(\tau-1)*d^{\text{stim}}}^{\tau*d^{\text{stim}}} g(t)$ to assign to $\beta_i^{\text{stim}}(\tau)$ (case A) or $\mu_i^{\text{stim}}(\tau)$ (case B).

Once all $\boldsymbol{\mu}_k$ have been set, $\boldsymbol{\beta}_i$ are sampled from $f(\boldsymbol{\beta}_i; \boldsymbol{\mu}_k, \sigma^2 * I)$, where they have not already been determined (case A). Once all of $\boldsymbol{\beta}_i$ has been determined, the spike train for the $i$th neuron is then simulated by sampling from the GLM's distribution for $y_i(1)$, then $y_i(2)$, all the way up to $y_i(T_i)$ (using eq 2).

### S5 Model Selection

#### S5.1 Alternative Model Selection: Validation Log-Likelihood

In addition to using BIC to perform model selection, we also investigate using the validation log-likelihood (VLL) on held out neurons (see section 2.4 detailed method).

For our simulated data, we use the exact same models that the BIC analysis was applied to (Figures 3,S2), but evaluate the model log-likelihood, averaged over a new validation set of 10 neurons per true cluster. Unlike with BIC, this VLL increases monotonically with $K$ towards an asymptote for both the simultaneous and sequential methods.

For each of the 50 simulated datasets, we selected the lowest $K$ whose VLL was within one standard error of the maximum, where this standard error was measured over the 50 datasets, after subtracting off each dataset's VLL for $K = 1$. This last step improves performance because each dataset has an offset in VLL that results from the spike density in the validation neurons (compare Figures S7 and S8 for an illustration of this effect in the IVSCC data). Overall, this approach can be thought of as a variant of the one-standard-error (1SE) rule [2]. These results are shown in Figure S6, and can be compared to those obtained via BIC in Figures 3,S2. For case A, BIC is better, but for case B, the results are more mixed. Across all cases, BIC tends to select a lower $\hat{K}$ than VLL.

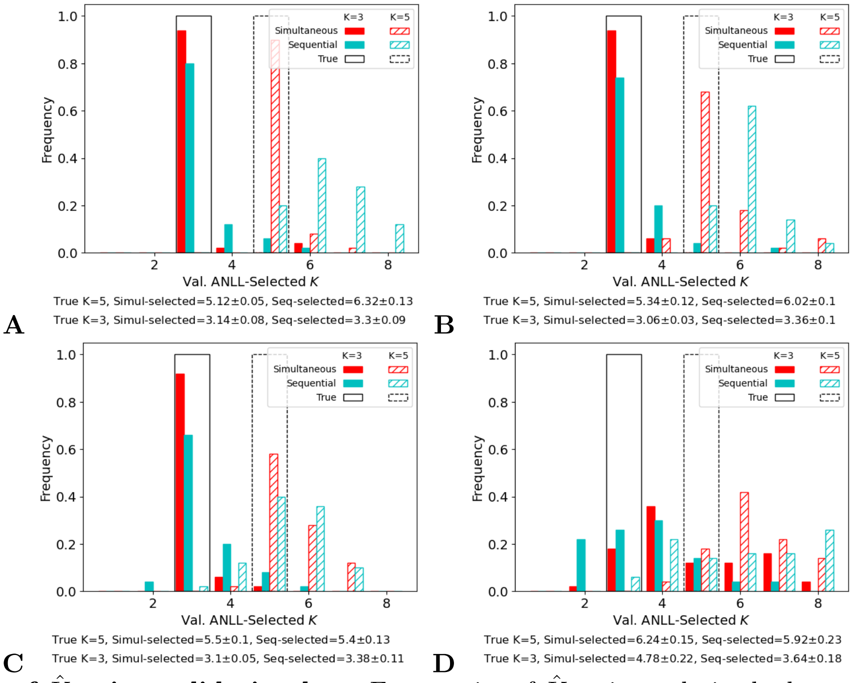


**Fig S6. Model selection of $\hat{K}$ using validation loss.** Frequencies of $\hat{K}$ estimated via the loss on held out neurons over 50 simulated datasets with the same $\mu_k$ as in Figure 2 (case A, panels A,B) or Figure S1 (case B, panels C,D), and $\sigma = 10^{-2}$ (A,C) or $10^{-5/6}$ (B,D), the maximum value that does not result in degenerate simulations. Black lines indicate true $K$. Summary below plots gives the mean ± SEM of estimated $\hat{K}$ across the 50 datasets for each case and each method.

For real data, where the true distribution of neurons is unknown, we must use cross-validated log-likelihood (CVLL) to get a metric that is not biased by our choice of test neurons. To compute CVLL, we randomly partition the neurons into $L$ equally-sized sets $A_1, ..., A_L$. Each set $A_l$ in turn is held out while the parameters are fit using $K$ clusters to the remaining neurons, and then the validation loss is simply $\frac{1}{|A_l|}\sum_{i \in A_l} \mathrm{LL}_i$ (see Section 2.4 for our definition of $\mathrm{LL}_i$). CVLL is then evaluated as the average validation loss over the $L$ different sets.

When this approach is applied to the IVSCC data, we observe the same issue with monotonic improvement in VLL with $K$ (Figure S7). Additionally, the simultaneous method yields VLLs for each validation fold that are offset far away from one another (Figure S7A,B). These offsets stem from large differences between the spike trains of individual neurons: some (with many spikes) yield a much higher ANLL, while others (with few spikes) yield a low ANLL (see Figures 5,S4 for an illustration of this phenomenon). Since the ANLL of all the validation cells factors into the validation loglikelihood (VLL) in the simultaneous method, a random partition of the neurons into different validation sets produces some sets with higher VLL and some with lower VLL.

To account for the offsets between these curves, we first shift them by subtracting off their values at $K = 1$ before using the standard 1SE rule to select $\hat{K}$ (Figure S8). That is, we pick the lowest $K$ whose CVLL is within one standard error of the maximum,

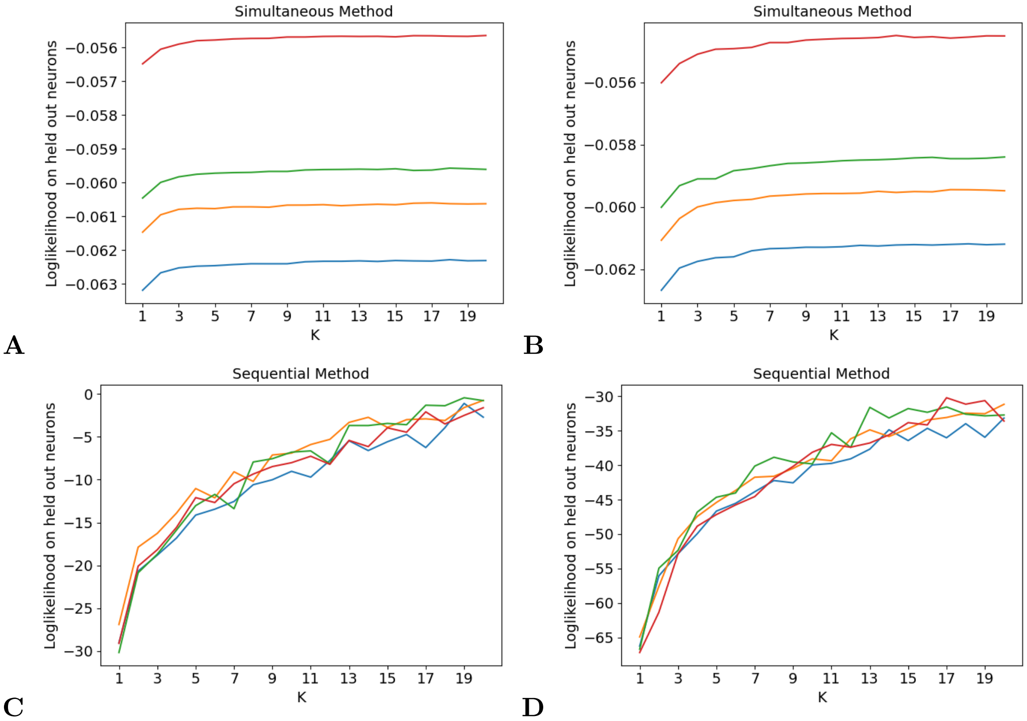


**Fig S7. Validation Log-Likelihood on IVSCC Dataset.** Each color represents a specific split of the neurons into training and testing. All log-likelihoods increase monotonically with $K$. The different folds of the simultaneous method are heavily offset relative to one another as a result of the variety of spike counts in each fold's validation set (see Figures 5,S4).
A: Case A, Simultaneous
B: Case B, Simultaneous
C: Case A, Sequential
D: Case B, Sequential

where the standard error is computed over the shifted VLLs of the different partitions of the neurons into training and validation.

Just as with the simulated datasets, CVLL selects a much higher $\hat{K}$ than BIC for each case (compare to Figure 4A,C and S3A,C). It is worth noting that as the VLL curves have not fully plateaued, repeating this analysis with a greater range of $K$ would likely result in a higher selected $\hat{K}$.

#### S5.2 Generalization Performance of Model Selection Metrics

We have now shown two different approaches to model selection and how they differ, both in method and results on simulated and real data. For the simulated data, as we know the true $K$, we are able to assess how the accuracy of selected $\hat{K}$. For the IVSCC data, we cannot do this directly as there is no ground truth for $K$, but we can attempt a similar approach to that used in Figures 5,S4, using the same measures of performance on predicting responses to the test stimulus of neurons that were held out from fitting the cluster models. Just as we argued that improvements in these metrics implied improved estimation of model parameters, $\boldsymbol{\beta}_i$, we can look for differences in these metrics, evaluated on test neurons, with respect to the $K$ used to fit the simultaneous method, and compare them to model selection criteria computed on the train/validation neurons. The $\boldsymbol{\beta}_i$ estimated by the sequential approach do not depend on $K$, so this analysis is limited to the sequential method.

However, these metrics do not show sufficient dependence on $K$ to warrant such an

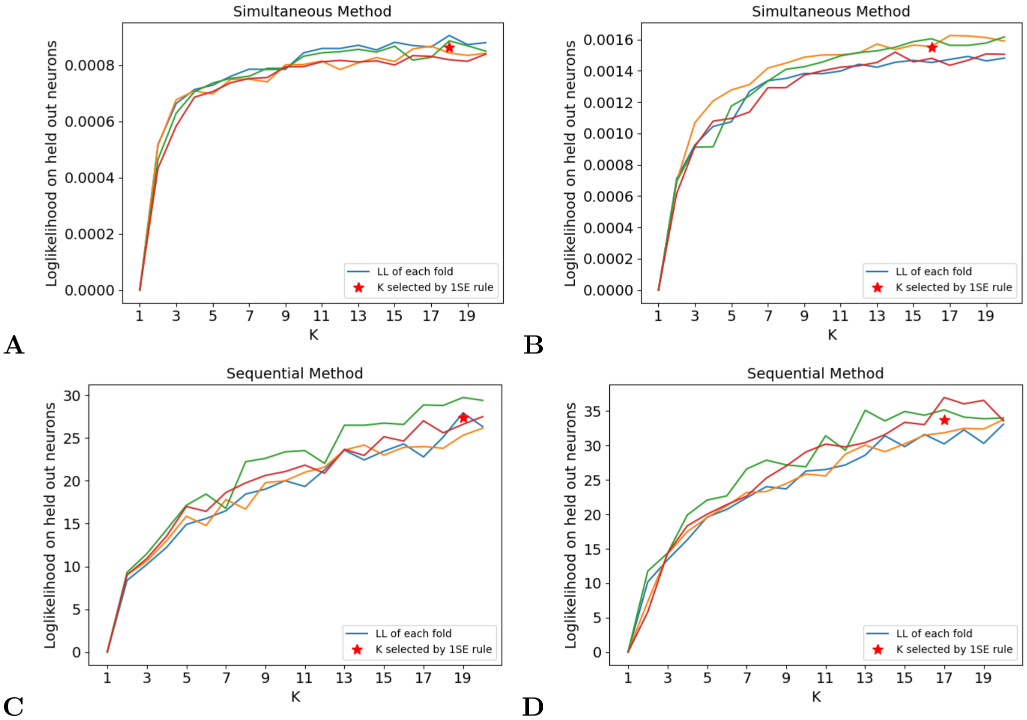


**Fig S8. Validation Log-Likelihood on IVSCC Dataset.** Same colors as Figure S7. The One-Standard-Error rule (1SE) applied to the Cross-Validated Loss (CVL) selects very high $\hat{K}$ for all methods (red stars). To account for the large offsets shown in Figure S7, each VL curve is first shifted (as shown here) so that its value for $K = 1$ is 0 before 1SE is applied.
A: Case A, Simultaneous
B: Case B, Simultaneous
C: Case A, Sequential
D: Case B, Sequential

analysis (Figure S9). With the possible exceptions of $K \in \{1, 2\}$, there is very little difference between the two cases, A and B, or between fitted $K$, relative to the variation between neurons.

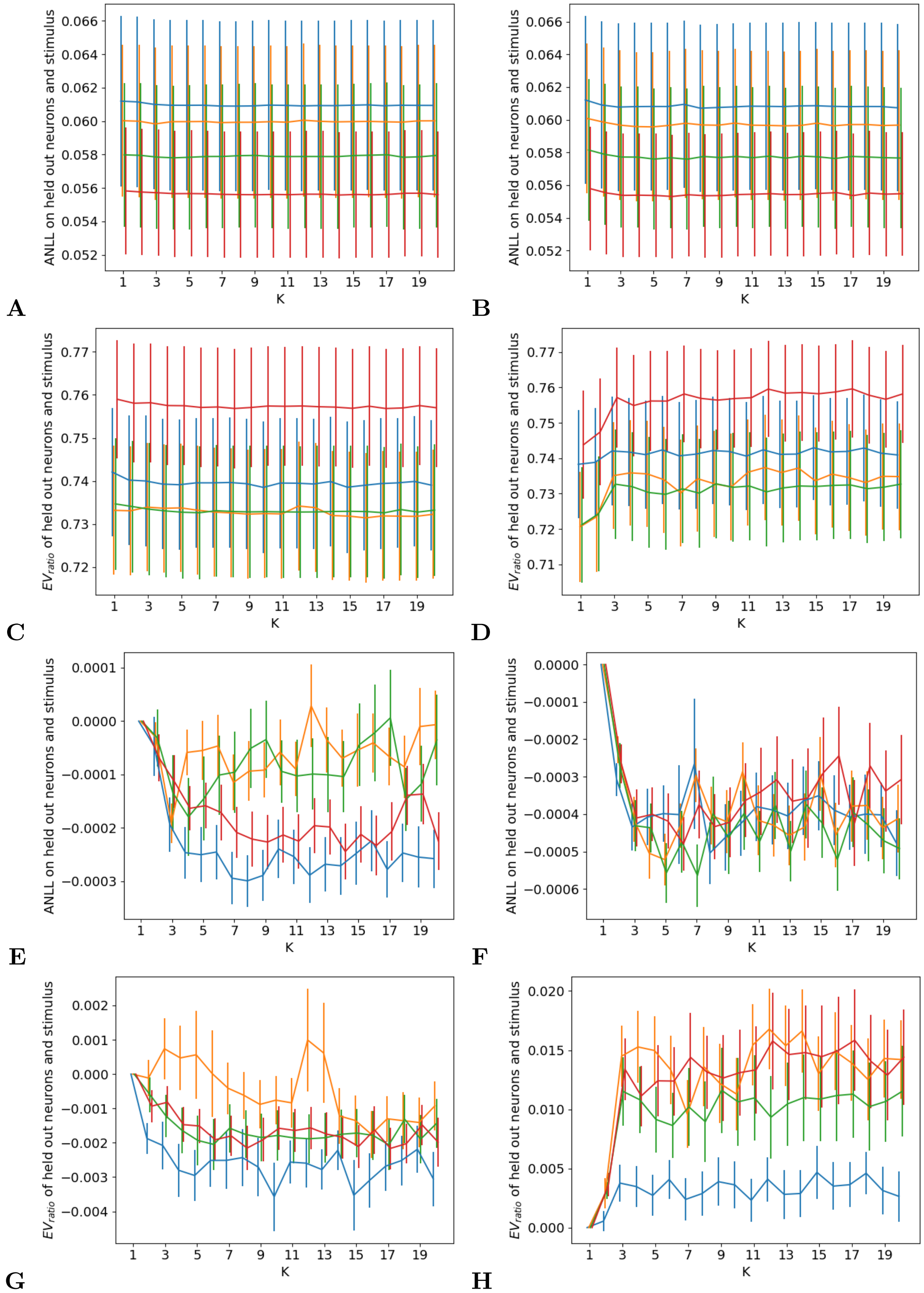


**Fig S9. Model Comparison using CVL on IVSCC Dataset.** Validation metrics for each fold are shown, with the same colors as in Figure S7. A-D: Error bars are one SE across neurons in the validation fold. E-H: The curves are shifted so that their values at $K = 1$ are 0, to facilitate judging their (lack of) change w.r.t. $K$. Error bars are one SE across validation neurons of the shifted metrics.
A,E: Case A, ANLL
B,F: Case B, ANLL
C,G: Case A, $EV_{ratio}$
D,H: Case B, $EV_{ratio}$
